# Putting focus on transcranial direct current stimulation in language production studies

**DOI:** 10.1101/230623

**Authors:** Jana Klaus, Dennis J.L.G. Schutter

## Abstract

**Objective:** Previous language production studies targeting the inferior frontal and superior temporal gyrus using anodal tDCS have provided mixed results. Part of this heterogeneity may be explained by limited target region focality of conventionally used electrode montages. We examined the focality of conventionally and alternative electrode montages.

**Methods:** Electrical field distributions of anodal tDCS targeting IFG and pSTG were simulated in conventional setups (anodal electrode over IFG/pSTG, reference electrode over right supraorbital region) and an alternative electrode montage in four different brains.

**Results:** Conventional montages showed maximum field strengths outside of the target regions. Results from alternative electrode montages showed that focality of tDCS could be improved by adjustments in electrode size and placement.

**Conclusions:** Heterogeneity of findings of language production studies deploying conventional tDCS montages may in part be explained by diffuse electrical field distributions. Alternative montages may improve focality and provide more unequivocal results.

**Significance:** Reliability of tDCS in language production research, both in basic and applied fields, can be improved by adopting different electrode montages which target the region of interest in a more direct way.

---

In studies on the functional neuroanatomy of language production transcranial direct current stimulation (tDCS) is routinely administered over the left inferior frontal gyrus (IFG) or posterior superior temporal gyrus (pSTG) in healthy volunteers. Results of these non-invasive brain stimulation studies have been subject to a significant degree of variability. For example, a number of studies reported a beneficial effect of anodal tDCS as evidenced by higher verbal fluency scores or shorter response times in picture naming tasks (Cattaneo, Pisoni, & Papagno, 2011; Fertonani, Rosini, Cotelli, Rossini, & Miniussi, 2010; Holland et al., 2011; Meinzer et al., 2012; Sparing, Dafotakis, Meister, Thirugnanasambandam, & Fink, 2008; Vannorsdall et al., 2012), while others did not find such an improvement (Cerruti & Schlaug, 2009; Ehlis, Haeussinger, Gastel, Fallgatter, & Plewnia, 2016; Henseler, Mädebach, Kotz, & Jescheniak, 2014; Vannorsdall et al., 2016; Westwood, Olson, Miall, Nappo, & Romani, 2017). Furthermore, meta-analyses on the ability of tDCS to modulate language performance in healthy volunteers have reported small or no effects (Horvath, Forte, & Carter, 2015; Klaus & Schutter, 2018; Price, McAdams, Grossman, & Hamilton, 2015; Westwood & Romani, 2017). Thus, even though tDCS may be effective in establishing effects on language processes, the heterogeneous results illustrate the difficulties associated with tDCS in anticipating both the direction and the magnitude of its behavioural effects (Jacobson, Koslowsky, & Lavidor, 2012; Oldrati & Schutter, 2017).

Issues concerning the spatial resolution of the induced electrical field are considered to be among the most important contributors to the diversity of tDCS-related effects. The vast majority of studies routinely placed one electrode over either the IFG or pSTG and the return electrode over the right supraorbital region (rSO, e.g. Cattaneo et al., 2011; Ehlis et al., 2016; Fiori, Cipollari, Caltagirone, & Marangolo, 2014; Pisoni, Cerciello, Cattaneo, & Papagno, 2017; Pisoni, Papagno, & Cattaneo, 2012; Westwood et al., 2017).^1^ Even though these montages have demonstrated the potential to be effective in manipulating processes underlying language production, computational simulation studies indicate that the intracranial electrical field distribution of tDCS is diffuse and the peak field strength amplitude is not located directly underneath the electrode (Rampersad et al., 2014). For instance, it has been shown that the maximum field strength between two electrodes is obtained if the target electrode is approximately placed between 20 and 40 mm away from the target region (Laakso et al., 2016; Rampersad et al., 2014). By this logic, placing the active electrode directly over the target site may induce a maximum field strength in regions adjacent to the targeted area. As a result, attributing changes in language production that are assumed to be caused by manipulating the regions of interest directly under the electrode can become more difficult. In spite of the well documented fact that electrode montage is an important aspect of tDCS experiments (Bikson, Datta, Rahman, & Scaturro, 2010; Laakso et al., 2016; Wagner et al., 2007), the extent to which suboptimal electrode montages may at least partially account for the heterogeneity of results in language production studies has not been examined yet. The goal of the present study was to (1) provide an estimate of the electrical field distributions of the two most commonly used electrode montages targeting the IFG and pSTG in language production tDCS studies, and (2) to present alternative ways to optimize the use of tDCS in language production studies.

## Material and methods

Using the SimNIBS software (version 2.0, Opitz, Paulus, Will, Antunes, & Thielscher, 2015) we simulated the electrical field distribution of anodal 1.5 mA tDCS with 5 × 7 cm electrodes (current density: 0.043 mA/cm^2^) in four different scenarios on four different brains. We used the T1-weighted resting-state structural magnetic resonance images from four participants provided in the publicly available dataset of the Sleepy Brain project (Nilsonne et al., 2016). We selected a male and female student from a young sample (between 20 and 30 years; participants 9001 and 9018 in the original dataset) and a male and female participant from an old sample (between 65 and 75 years; participants 9002 and 9004). We first created individual tetrahedral head models using the mri2mesh algorithm implemented in the SimNIBS pipeline (see Windhoff, Opitz, & Thielscher, 2013, for a more specific description of the procedure). Then we tested whether the conventionally used montage of placing the active electrode over the target region and the reference electrode over the contralateral supraorbital region provides the desired focality across the left IFG and pSTG, respectively. To address the second aim of our study, we used the computational findings reported by Rampersad et al. (2014) to adjust the electrode positions in the following ways (1) moving the centre of the active electrode approximately 3 cm anterior to the target region; (2) placing the reference electrode in closer proximity to the active electrode; (3) turning the electrode so that the short edges of both electrodes approximately face each other. Electrode size was kept constant in order to directly compare the influence of electrode placement on the focality of tDCS. For all simulations, we chose an electrode-sponge setup, with a 1 mm thick electrode covered by a 2 mm thick sponge.

## Results

Table 1 reports the individual minimum and maximum electrical field strengths (in V/m) broken down by stimulation site (IFG vs. pSTG) and montage (conventional vs. alternative). Note, however, that these values refer to field strengths observed *anywhere* in the brain and do not speak to the question which electrical fields are elicited in the target regions. Nevertheless, it does illustrate both inter- and intraindividual variability in the magnitude of the generated electrical field, as has also been reported in a large sample by Laakso et al. (2016).

**Table 1.**
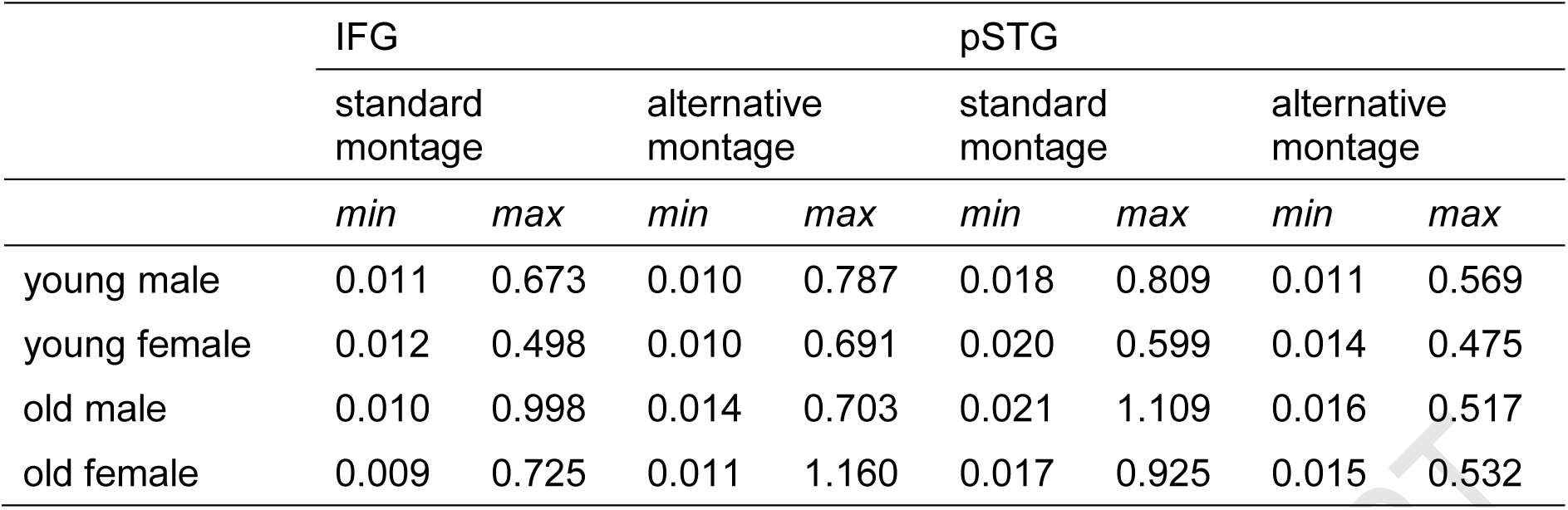
Minimum and maximum electrical field strengths (in V/m) for all simulations.

### IFG montage

Figure 1 displays the electrode montages and simulation results for the montages targeting the left IFG in the four brains. Results indicate that the conventionally used montage (left panel of Figure 1) is not optimal as peaks in field strength were found in bilateral middle and superior frontal regions. Furthermore, the field distributions spread with decreasing intensity to left central and anterior temporal regions. Thus, while the standard montage did affect the target region, substantially higher electrical fields were observed anterior to the region of interest, and in both hemispheres.

**Figure 1.**
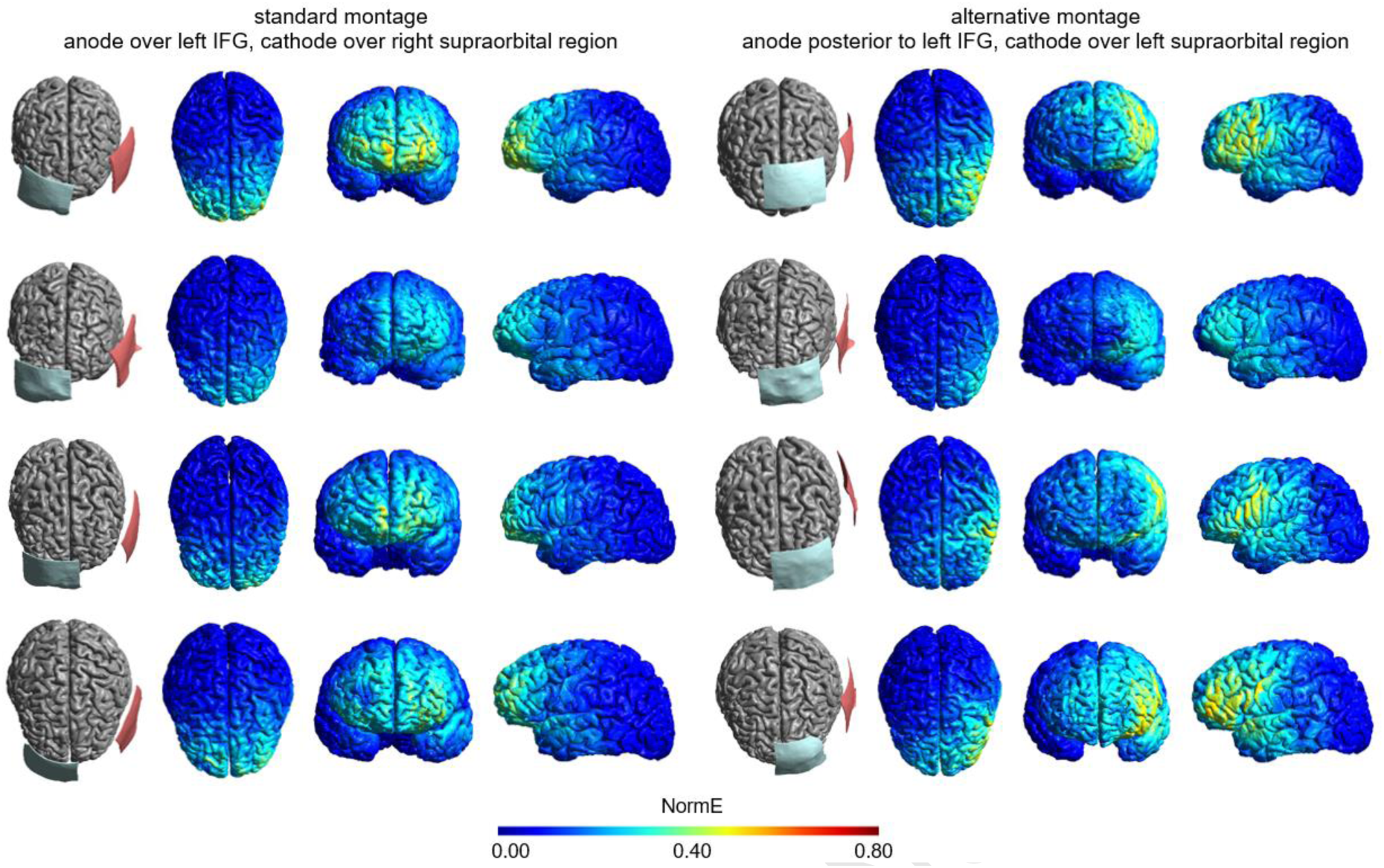
Electrode montages and electrical field intensities for tDCS targeting the left inferior frontal gyrus for the four participants (from top to bottom: young male, young female, old male, old female). The left part displays the simulation results for the conventional montage in which the anodal electrode is placed over the left IFG and the cathodal electrode is placed over the right supraorbital region. The right part displays the simulation results for the alternative montage in which the anodal electrode is placed posterior to the left IFG and the cathodal electrode is placed over the left supraorbital region. All electrical fields are scaled between 0 and 0.8 V/m, with brighter colours indicating higher electrical field strengths.

The alternative electrode montage (right panel of Figure 1) showed a notable shift of the electrical field. While the magnitude and spread of the induced electrical field differed between individuals, the simulations uniformly displayed a convergence to the target area. That is, the effect on the right hemisphere was reduced, and the peak intensities were located around the IFG and the central sulcus, with additional, decreased field intensities in middle frontal and anterior temporal regions.

### pSTG montage

Figure 2 shows the results from simulations targeting the pSTG. The conventional montage (left part of Figure 2) resulted in a wide electrical field distribution in the left hemisphere. Field strength peaks were centred on the medial part of the postcentral gyrus (i.e., anterior to the target region). Additionally, the induced electrical field covered significant parts of the left hemisphere and anterior parts of the contralateral hemisphere, arguably due to the large distance between the electrodes. As for IFG stimulation, the alternative montage (right panel of Figure 2) shifted the field intensity peaks towards the target region, with the highest field strength found in a rather large region including the pSTG and the inferior parietal lobule. Right-hemispheric effects were eliminated entirely because for pSTG stimulation, the alternative montage exclusively placed the electrodes across the left hemisphere, thus preventing the generation of electric fields in the right hemisphere.

**Figure 2.**
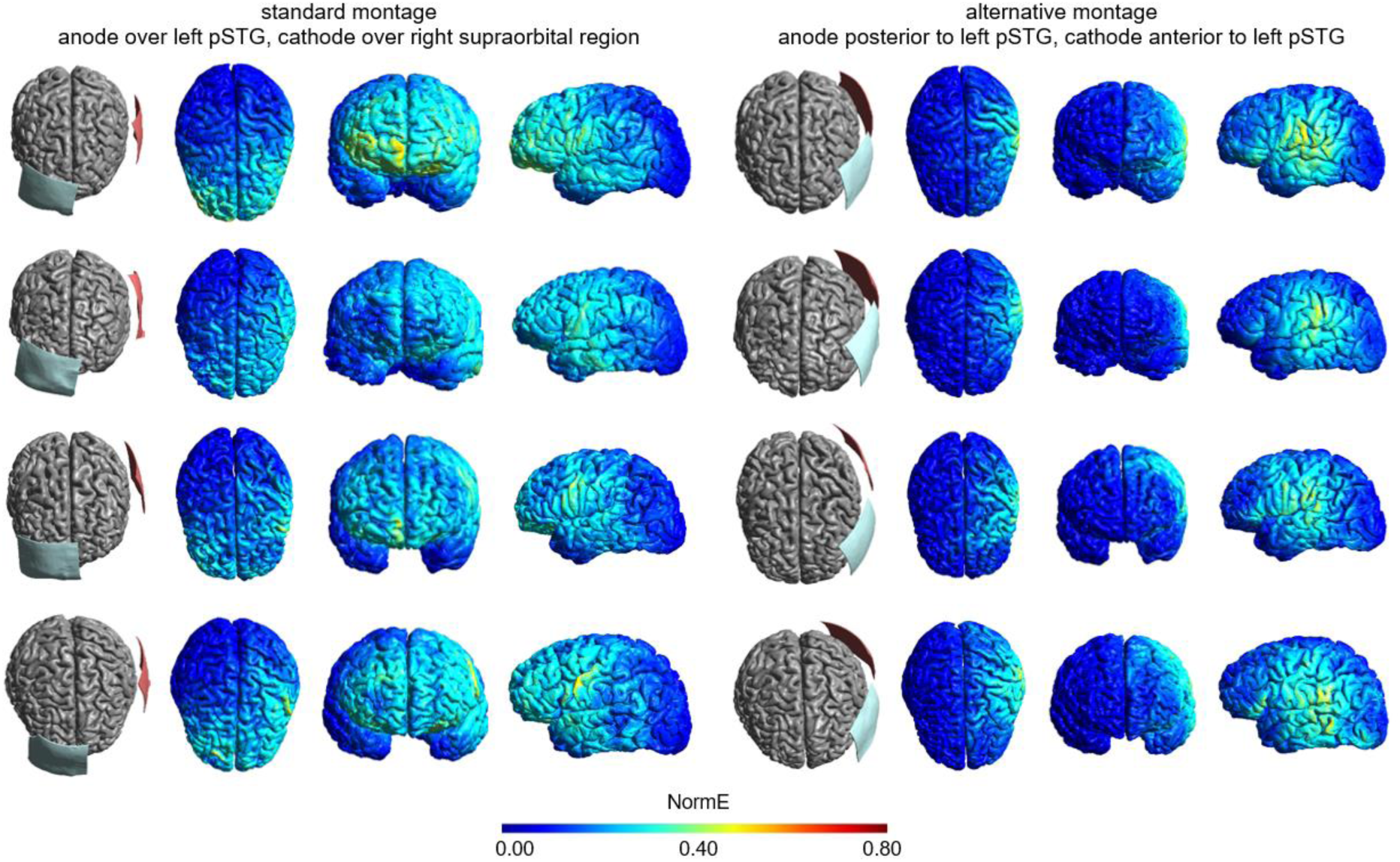
Electrode montages and electrical field intensities for tDCS targeting the left posterior superior temporal gyrus for the four participants (from top to bottom: young male, young female, old male, old female). The left part displays the simulation results for the conventional montage in which the anodal electrode is placed over the left pSTG and the cathodal electrode is placed over the right supraorbital region. The right part displays the simulation results for the alternative montage in which the anodal electrode is placed posterior to the left pSTG and the cathodal electrode is placed anterior to the left pSTG. All electrical fields are scaled between 0 and 0.8 V/m, with brighter colours indicating higher electrical field strengths.

## Discussion

Our simulation results showed that the electrical field distribution of tDCS montages most commonly used in language production studies can be improved. Placement of the active electrode over the target region and the reference electrode over the contralateral supraorbital region yields the highest field strengths anterior to the target region as well as additional frontal effects in the right hemisphere. These wide electrical field distributions may cause collateral activation of surrounding tissue and contribute to the heterogeneous findings reported in previous studies. While there is no immediate reason to assume that the target regions were not exposed to the exogenous electrical field at all in previous studies, conventionally applied montages may have been suboptimal in reaching the desired spatial resolution of the electrical field. Consequently, small effects of tDCS on language production may have been caused, at least in part, by affecting the target region to varying degrees.

Here, we provide an alternative montage that based on computer simulations produced more focal peaks in electrical field strength. Altering the montage by placing the electrodes anterior and posterior to the target region improved focality in all of our four brains. Using such a montage may thus target the desired region more directly and limit electrical field exposure of surrounding regions. As a result, future studies reporting differences between real and sham tDCS conditions may find more reliable and unequivocal effects in language production tasks when targeting a region of interest. The magnitude of the induced electrical field appears to be well in the range required to elicit cellular effects (Rahman, Lafon, & Bikson, 2015). Nevertheless, it remains to be tested whether the improved montages are indeed effective in obtained less unequivocal effects of tDCS in language production performance. Additionally, as can be seen from the simulations, the cortical area affected by the stimulation typically covers a large region between the electrodes. It remains to be tested whether additional modifications to the montages (e.g. by using smaller electrodes or a high-definition tDCS setup) further reduce induced field strengths in regions peripheral to the target region.

Aside from the basic neuroscientific questions on language production, tDCS is currently being explored as a possible therapeutic intervention to treat aphasia (Elsner, Kugler, Pohl, & Mehrholz, 2015; Sandars, Cloutman, & Woollams, 2016; Sebastian, Tsapkini, & Tippett, 2016). Based on our findings, it is reasonable to assume that aphasic patients treated with tDCS may benefit from adopting simulation-based electrode montages to maximize focal electrical field strengths (see also Galletta et al., 2015, for a simulation-based approach in a lesioned brain).

It should be noted that the simulations were performed on a small number of brains, which limits the generalisability of our results. Evidently, the efficacy of tDCS depends on many factors that include individual differences in gyral folding and physiological susceptibility to weak electrical currents—a source of variability which is also visible in the current simulations. Nevertheless, we have demonstrated that despite alleged individual differences in electrical field distributions, the alternative montages were consistently more successful in targeting the desired regions. Thus, irrespective of anatomical and cellular differences between participants, our simulations provide evidence that the standard montage is suboptimal. Finally, research labs and clinics that do not have direct access to neuroimaging facilities to individualize tDCS montages, simulations provide a pragmatic solution that outweighs the alternative of relying on conventionally-used montages.

In conclusion, our results show that the type of montage may contribute to the robustness of findings, improve their interpretations, and advance the application of tDCS on language production performance in basic neuroscientific and clinical settings.

## Acknowledgments

This work was supported by the German Research Council under Grant KL 2933/2.

## Footnotes

1 Note that some studies also used the vertex, contralateral homologue, or extracephalic locations (e.g., contralateral cheek or shoulder) as reference positions (Sparing et al., 2008; Vannorsdall et al., 2012; Westwood et al., 2017; Wirth et al., 2011). Although we here focus on the rSO as the reference region, additional simulations (not reported here) including the vertex or the contralateral homologue provided comparable results (i.e., highest field strengths outside of the targeted region).

## References

Bikson, M., Datta, A., Rahman, A., & Scaturro, J. (2010). Electrode montages for tDCS and weak transcranial electrical stimulation: role of “return” electrode’s position and size. Clinical Neurophysiology, 121(12), 1976–8.

Cattaneo, Z., Pisoni, A., & Papagno, C. (2011). Transcranial direct current stimulation over Broca’s region improves phonemic and semantic fluency in healthy individuals. Neuroscience, 183, 64–70.

Cerruti, C., & Schlaug, G. (2009). Anodal transcranial direct current stimulation of the prefrontal cortex enhances complex verbal associative thought. Journal of Cognitive Neuroscience, 21(10), 1980–7.

Ehlis, A.-C., Haeussinger, F. B., Gastel, A., Fallgatter, A. J., & Plewnia, C. (2016). Task-dependent and polarity-specific effects of prefrontal transcranial direct current stimulation on cortical activation during word fluency. NeuroImage, 140, 134–140.

Elsner, B., Kugler, J., Pohl, M., & Mehrholz, J. (2015). Transcranial direct current stimulation (tDCS) for improving aphasia in patients with aphasia after stroke. Cochrane Database of Systematic Reviews, 25(6), CD009760.

Fertonani, A., Rosini, S., Cotelli, M., Rossini, P. M., & Miniussi, C. (2010). Naming facilitation induced by transcranial direct current stimulation. Behavioural Brain Research, 208(2), 311–318.

Fiori, V., Cipollari, S., Caltagirone, C., & Marangolo, P. (2014). “If two witches would watch two watches, which witch would watch which watch?” tDCS over the left frontal region modulates tongue twister repetition in healthy subjects. Neuroscience, 256, 195–200.

Galletta, E. E., Cancelli, A., Cottone, C., Simonelli, I., Tecchio, F., Bikson, M., & Marangolo, P. (2015). Use of computational modeling to inform tDCS electrode montages for the promotion of language recovery in post-stroke aphasia. Brain Stimulation, 8(6), 1108–1115.

Henseler, I., Mädebach, A., Kotz, S. A., & Jescheniak, J. D. (2014). Modulating brain mechanisms resolving lexico-semantic interference during word production: A transcranial direct current stimulation study. Journal of Cognitive Neuroscience, 26(7), 1403–1417.

Holland, R., Leff, A. P., Josephs, O., Galea, J. M., Desikan, M., Price, C. J., et al. (2011). Speech facilitation by left inferior frontal cortex stimulation. Current Biology, 21(16), 1403–1407.

Horvath, J. C., Forte, J. D., & Carter, O. (2015). Quantitative review finds no evidence of cognitive effects in healthy populations from single-session transcranial direct current stimulation (tDCS). Brain Stimulation, 8(3), 535–550.

Jacobson, L., Koslowsky, M., & Lavidor, M. (2012). tDCS polarity effects in motor and cognitive domains: a meta-analytical review. Experimental Brain Research, 216(1), 1–10.

Klaus, J., & Schutter, D. J. L. G. (2018). Non-invasive brain stimulation to investigate language production in healthy speakers: A meta-analysis. Brain and Cognition, 123.

Laakso, I., Tanaka, S., Mikkonen, M., Koyama, S., Sadato, N., & Hirata, A. (2016). Electric fields of motor and frontal tDCS in a standard brain space: A computer simulation study. NeuroImage, 137, 140–151.

Meinzer, M., Antonenko, D., Lindenberg, R., Hetzer, S., Ulm, L., Avirame, K., et al. (2012). Electrical brain stimulation improves cognitive performance by modulating functional connectivity and task-specific activation. The Journal of Neuroscience, 32(5), 1859–1866.

Nilsonne, G., Tamm, S., d’Onofrio, P., Thuné, H. Å., Schwarz, J., Lavebratt, C., et al. (2016). A multimodal brain imaging dataset on sleep deprivation in young and old humans. Retrieved from https:openarchive.ki.se/xmlui/handle/10616/45181

Oldrati, V., & Schutter, D. J. L. G. (2017). Targeting the human cerebellum with transcranial direct current stimulation to modulate behavior: A meta-analysis. The Cerebellum, 1–9.

Opitz, A., Paulus, W., Will, S., Antunes, A., & Thielscher, A. (2015). Determinants of the electric field during transcranial direct current stimulation. NeuroImage, 109, 140–150.

Pisoni, A., Cerciello, M., Cattaneo, Z., & Papagno, C. (2017). Phonological facilitation in picture naming: When and where? A tDCS study. Neuroscience, 352, 106–121.

Pisoni, A., Papagno, C., & Cattaneo, Z. (2012). Neural correlates of the semantic interference effect: New evidence from transcranial direct current stimulation. Neuroscience, 223, 56–67.

Price, A. R., McAdams, H., Grossman, M., & Hamilton, R. H. (2015). A meta-analysis of transcranial direct current stimulation studies examining the reliability of effects on language measures. Brain Stimulation, 8(6), 1093–1100.

Rahman, A., Lafon, B., & Bikson, M. (2015). Multilevel computational models for predicting the cellular effects of noninvasive brain stimulation. Progress in Brain Research, 222, 25–40.

Rampersad, S. M., Janssen, A. M., Lucka, F., Aydin, U., Lanfer, B., Lew, S., et al. (2014). Simulating transcranial direct current stimulation with a detailed anisotropic human head model. IEEE Transactions on Neural Systems and Rehabilitation Engineering, 22(3), 441–452.

Sandars, M., Cloutman, L., & Woollams, A. M. (2016). Taking sides: An integrative review of the impact of laterality and polarity on efficacy of therapeutic transcranial direct current stimulation for anomia in chronic poststroke aphasia. Neural Plasticity, 2016, 1–21.

Sebastian, R., Tsapkini, K., & Tippett, D. C. (2016). Transcranial direct current stimulation in post stroke aphasia and primary progressive aphasia: Current knowledge and future clinical applications. NeuroRehabilitation, 39(1), 141–152.

Sparing, R., Dafotakis, M., Meister, I. G., Thirugnanasambandam, N., & Fink, G. R. (2008). Enhancing language performance with non-invasive brain stimulation—A transcranial direct current stimulation study in healthy humans. Neuropsychologia, 46(1), 261–268.

Vannorsdall, T. D., Schretlen, D. J., Andrejczuk, M., Ledoux, K., Bosley, L. V, Weaver, J. R., et al. (2012). Altering automatic verbal processes with transcranial direct current stimulation. Frontiers in Psychiatry, 3, 73.

Vannorsdall, T. D., van Steenburgh, J. J., Schretlen, D. J., Jayatillake, R., Skolasky, R. L., & Gordon, B. (2016). Reproducibility of tDCS results in a randomized trial. Cognitive And Behavioral Neurology, 29(1), 11–17.

Wagner, T., Fregni, F., Fecteau, S., Grodzinsky, A., Zahn, M., & Pascual-Leone, A. (2007). Transcranial direct current stimulation: A computer-based human model study. NeuroImage, 35(3), 1113–1124.

Westwood, S. J., Olson, A., Miall, R. C., Nappo, R., & Romani, C. (2017). Limits to tDCS effects in language: Failures to modulate word production in healthy participants with frontal or temporal tDCS. Cortex, 86, 64–82. http:doi.org/10.1016/j.cortex.2016.10.016

Westwood, S. J., & Romani, C. (2017). Transcranial direct current stimulation (tDCS) modulation of picture naming and word reading: A meta-analysis of single session tDCS applied to healthy participants. Neuropsychologia.

Windhoff, M., Opitz, A., & Thielscher, A. (2013). Electric field calculations in brain stimulation based on finite elements: An optimized processing pipeline for the generation and usage of accurate individual head models. Human Brain Mapping, 34(4), 923–935.

Wirth, M., Rahman, R. A., Kuenecke, J., Koenig, T., Horn, H., Sommer, W., & Dierks, T. (2011). Effects of transcranial direct current stimulation (tDCS) on behaviour and electrophysiology of language production. Neuropsychologia, 49(14), 3989–3998. http:doi.org/10.1016/j.neuropsychologia.2011.10.015

